# BioBrain: A Multi-Agent Framework for Natural Language Driven Quantitative Microscopy Data Analysis

**DOI:** 10.64898/2026.06.17.732700

**Authors:** K. Tsolakidis, Artu Breuer, S. Bender, Stavroula Margaritaki, Marcus W. Dreisler, A. Oikonomou, Nikos S Hatzakis

## Abstract

Advances in fluorescence microscopy have dramatically expanded the range of biological questions that can be addressed, enabling quantitative observations of molecular interactions and cellular dynamics with unprecedented spatial and temporal resolution. However, the growing complexity of imaging data has outpaced our ability to analyze them. Despite numerous computational methods exist, they often rely on specialized software environments, heterogeneous data formats, and technical expertise, limiting adoption and widening the gap between data acquisition and quantitative biological interpretation. Here we introduce BioBrain, a multi-agent framework that translates natural-language analytical goals into executable and reproducible microscopy analysis pipelines. Instead of generating analysis code, BioBrain assembles validated analytical methods and can expands its analytical capabilities by integrating existing laboratory scripts into a unified conversational framework. Every selected method and inferred parameter is transparently reported, ensuring traceable and reproducible analyses. On two-channel total internal reflection fluorescence and three-dimensional lattice light-sheet benchmarks, BioBrain exactly reproduces expert-derived results when parameters are specified and degrades predictably and traceably when they are not, while frontier language models generated large, model-dependent quantitative errors despite completing without warning. BioBrain offers a practical path for closing the widening gap between data acquisition and biological discovery, enabling experimental scientists to communicate with computational analysis in the language of biology rather than the language of software.

## Introduction

Advances in fluorescence microscopy have transformed our ability to investigate biology at the molecular and cellular scale. Developments such as super-resolution imaging, lattice light-sheet microscopy, and high-speed live-cell imaging now enable quantitative observation of molecular interactions, intracellular trafficking, and organelle dynamics with unprecedented spatial and temporal resolution. As a result, researchers can now ask quantitative biological questions that were largely inaccessible only a decade ago, for example, how many nanoparticles enter an individual cell, which intracellular pathways they follow, and when and where cargo is released. Consequently, the principal bottleneck has shifted from data acquisition to data analysis and interpretation^1^. Answering these biological questions requires utilizing complex computational workflows that typically involve heterogeneous software tools, custom scripts, and extensive parameter optimization. A scientist who has acquired a multi-gigabyte imaging dataset must often parse instrument-specific image formats, correct background signal, define segmentation or detection thresholds, identify cells or particles, extract quantitative fluorescence features, and integrate outputs across software tools that rarely share common interfaces. Each step demands both algorithmic and software-engineering expertise, a combination few experimentalists have alongside their primary research domain^2–4^.

Several barriers currently stand between researchers and quantitative biological insight, which can be broadly categorized into two types: Established graphical platforms such as ImageJ/Fiji, CellProfiler, and Napari lower the barrier to running individual algorithms but still require users to know which methods to use, tune parameters, and chain processing steps by hand^5–7^. Genera-purpose workflow engines such as Nextflow, Galaxy, and KNIME provide modular pipeline execution but must be configured explicitly and are not tailored to microscopy-specific tasks^8–10^. Meanwhile, the community continuously develops state-of-the-art computational methods spanning particle tracking, image reconstruction, cell tracking, large-scale light-sheet analysis, and foundation-model-based segmentation^11–18^. In principle, these methods while publicly available in practice, often remain effectively stranded between the literature and the bench. Accessing them demands navigating heterogeneous input formats, software dependencies, and execution environments that reflect the genuine complexity of the underlying analyses, a barrier that falls disproportionately on experimentalists rather than on the developers who built them.^19^. Ironically, as computational microscopy has become more powerful, it has also become less accessible to the experimental scientists who generated the data.

Recent advances in large language models (LLMs) suggest a solution: natural-language interfaces that translate experimental intent into executable analysis. Existing LLM-based systems for bioimage analysis largely fall into two categories. Conversational assistants such as the BioImage.IO Chatbot and Omega use retrieval-augmented generation and code synthesis to help users navigate tool ecosystems or perform individual processing steps^20,21^, but they operate in an advisory or single-step execution mode, and do not autonomously assemble multi-step workflows. General-purpose LLM code-generation agents can attempt to produce full analysis scripts from natural-language prompts^22,23^, but without domain constraints and schema-enforced interfaces, they can select inappropriate methods, infer incorrect parameters, or compose logically inconsistent processing steps^24–26^. Because such scripts typically execute without explicit failure, the resulting errors remain silent, and yield plausible-looking but incorrect quantitative outputs^27^. Together, these limitations define a clear unmet need for a fundamentally different paradigm: a system that translates natural-language descriptions of biological questions into reproducible workflows while designed to minimize silent analytical failures, one that is constrained to validated analytical methods, halts to request clarification from the user when intent is ambiguous, and exposes every analytical decision, making the process traceable and transparent.

To address this challenge, we developed BioBrain, a multi-agent LLM-powered framework that translates natural-language descriptions of biological questions into executable and reproducible microscopy analysis workflows. The coordinating Orchestrator Agent interprets the user’s natural-language request and decomposes it into subtasks that are delegated to specialized Sub-Agents responsible for specific analytical domains, which select appropriate tools from curated libraries of deterministic, schema-constrained functions. The modular architecture allows multistep workflows to be dynamically assembled while maintaining flexibility when new workflows are designed.

Three key design principles separate BioBrain from existing approaches. First, reasoning is separated from execution: the LLM plans the analysis, but execution is restricted to validated analytical tools; if the prompt is ambiguous or insufficiently specified, the framework requests clarification rather than making unsupported assumptions, and every selected tool, inferred parameter, and analytical decision is reported to ensure transparency and reproducibility. Second, raw microscopy data are never exposed to the language model, which operates only on user prompts and tool descriptions. Third, the analytical toolset is extensible through automated integration of existing laboratory scripts, so compatible laboratory-developed analysis methods can be incorporated without dedicated data-science effort. We validate BioBrain against pure LLM-based execution and on two representative microscopy analyses, total internal reflection fluorescence (TIRF) colocalization quantifying cargo release over time and three-dimensional lattice light-sheet quantification of nanoparticle internalization, showing that it exactly reproduces expert-derived results when parameters are specified and degrades predictably and traceably when they are not. By reducing the need for computational specialists, BioBrain makes advanced microscopy analysis as accessible as describing an experiment, without compromising the methodological standards that quantitative science requires.

## 3 Results

### From natural language to quantitative microscopy results

BioBrain enables experimental scientists to obtain quantitative measurements from microscopy data by expressing analytical objectives in natural language rather than computational terms. The user provides two inputs: a microscopy dataset in a supported format and a description of the analytical objective, such as quantifying time-resolved cargo-membrane colocalization or measuring particle internalization in a three-dimensional cell volume. BioBrain automatically translates this objective into an executable analysis workflow, performs the analysis, and returns the quantitative results together with a transparent record of the analytical process, including the selected tools, parameter values, and the origin of each parameter (Fig. 1a-c).

**Fig. 1.**
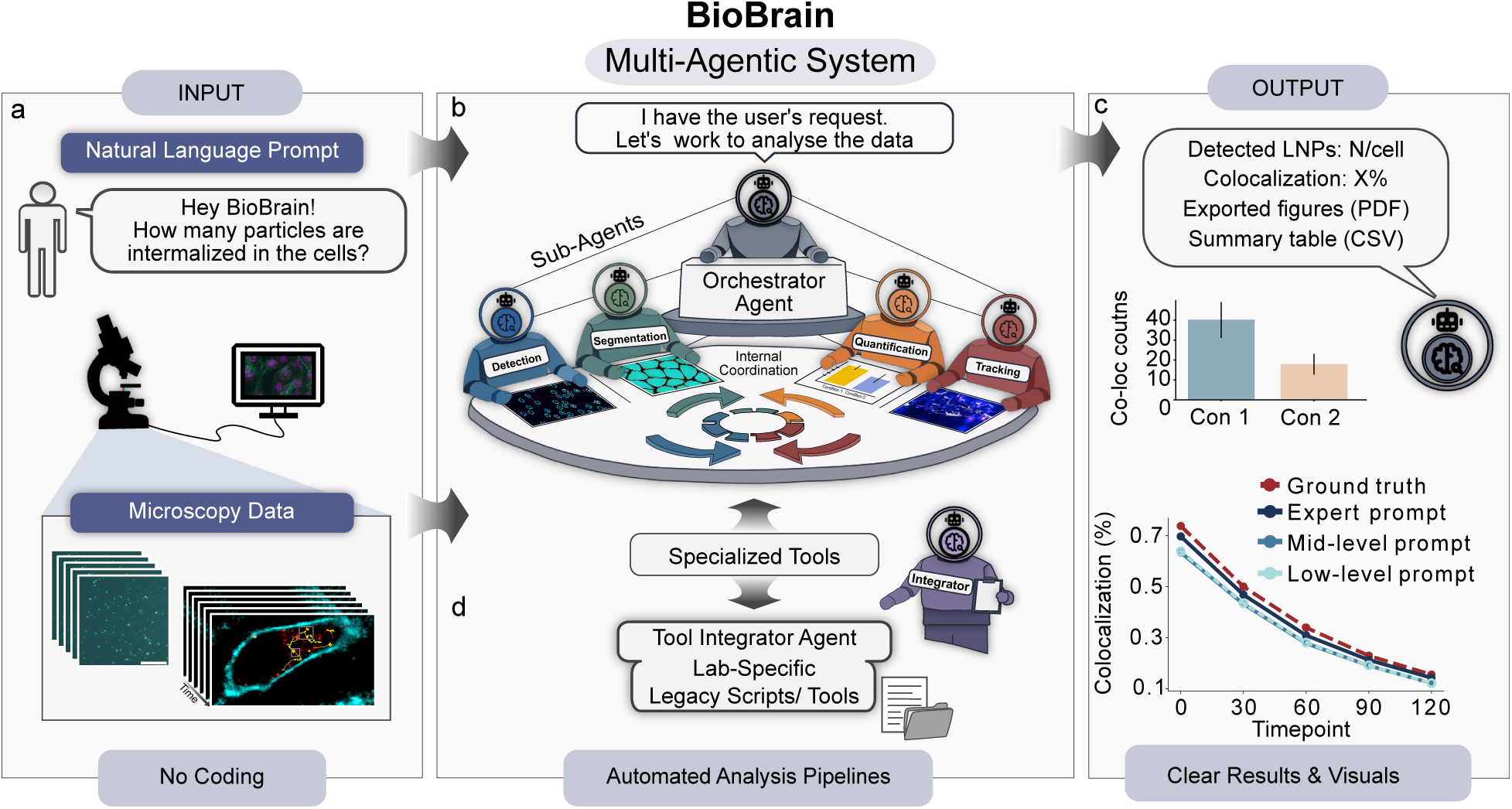
| BioBrain, a multi-agent LLM framework enabling natural-language-driven microscopy analysis. Schematic overview of the BioBrain system architecture and workflow. (a) A scientist provides a natural-language description of their analytical goal together with a path to the raw microscopy data in any supported format. (b) The Orchestrator Agent interprets the request, decomposes it into discrete computational subtasks, and dynamically assembles an executable workflow by coordinating domain-specific Sub-Agents, each restricted to a curated toolset. If required information is missing or an appropriate workflow cannot be identified with sufficient confidence, execution halts and clarification is requested before proceeding. (c) Results are returned to the user alongside a natural-language explanation, making outputs transparent and interpretable without specialist knowledge. (d) The Tool Integrator Agent extends the system’s analytical repertoire by automatically converting existing laboratory scripts into standardised, BioBrain-compatible tools, enabling integration of new methods without data-science intervention.

The level of user specification can vary from explicit algorithm choices and numerical parameters to qualitative descriptions or simply the underlying biological objective. When essential information is missing or ambiguous, BioBrain returns control to the user for clarification rather than proceeding under uncertainty, a behaviour that follows from the layered architecture described below.

BioBrain achieves this functionality through a layered architecture that separates reasoning, workflow planning, and deterministic execution (Supplementary Figs. S1, S2). A coordinating orchestrator agent interprets the user’s request and decomposes the analytical objective into a sequence of subtasks. These are delegated to specialized agents responsible for domains such as detection, segmentation, colocalization, and microscopy file handling. The computational work is performed by an underlying layer of deterministic Python tools exposed through Model Context Protocol (MCP) servers^28^. MCP provides a uniform interface through which tools can be registered with explicit descriptions and typed input-output schemas, allowing analytical functions to be invoked consistently across workflows. Consequently, the LLM plans the analysis and selects among registered analytical methods but execution remains restricted to validated tool calls rather than unconstrained code generation.

This separation enables one of BioBrain’s central design principles: transparent and constrained workflow generation. Rather than attempting to complete every request, BioBrain is designed to halt and request clarification, if essential information is missing, the requested analysis is ambiguous, or no suitable validated method can be identified (Supplementary Fig. S3). Designing the language model to only invoke registered analytical tools, rather than generate executable code, practically eliminates code-level hallucinations, such as inventing functions, methods, or API calls. When users provide qualitative prompts rather than explicit numerical parameters, BioBrain infers parameter values from the stated biological objective. These inferred values remain constrained to valid ranges supported by the selected tools. Every selected tool and inferred parameter is automatically recorded and reported, ensuring that the complete analytical workflow remains transparent, inspectable and reproducible, minimizing silent errors.

The same architecture also limits the information exposed to the language model. During execution, the LLM receives only the user’s natural-language request and descriptions of the available analytical tools; raw microscopy data and intermediate analysis outputs remain accessible only to the underlying analysis tools that perform the analysis. Thus, BioBrain functions as a natural-language interface in which AI guides workflow construction while quantitative analysis remains grounded in validated methods, transparent decision-making, and reproducible execution.

### Automated tool integration

A large portion of microscopy analysis pipelines exists not as reusable software packages but as laboratory-specific or published Python scripts. Although these scripts often implement state-of-the-art analyses, adopting them in a new laboratory typically requires substantial computational expertise to understand their dependencies, expected inputs and outputs, and execution environment. Consequently, many published methods remain effectively inaccessible despite being publicly available.

BioBrain addresses this barrier through a Tool Integrator Agent, which converts an existing script, once, into a BioBrain-compatible tool that can subsequently be selected by sub-agents and invoked through natural language (Fig. 2). To add a new method to BioBrain’s analytical toolbox, the user provides the Tool Integrator Agent with a Python script and a short description of the script’s function. The Tool Integrator Agent inspects the code using programmatic Abstract Syntax Tree (AST) analysis to identify public functions, imports, dependencies, input-output behaviour, and generates a standardised Model Context Protocol (MCP) wrapper that exposes the selected function with explicit typed input-output schemas and registers the resulting tool in the BioBrain tool layer.

**Fig. 2.**
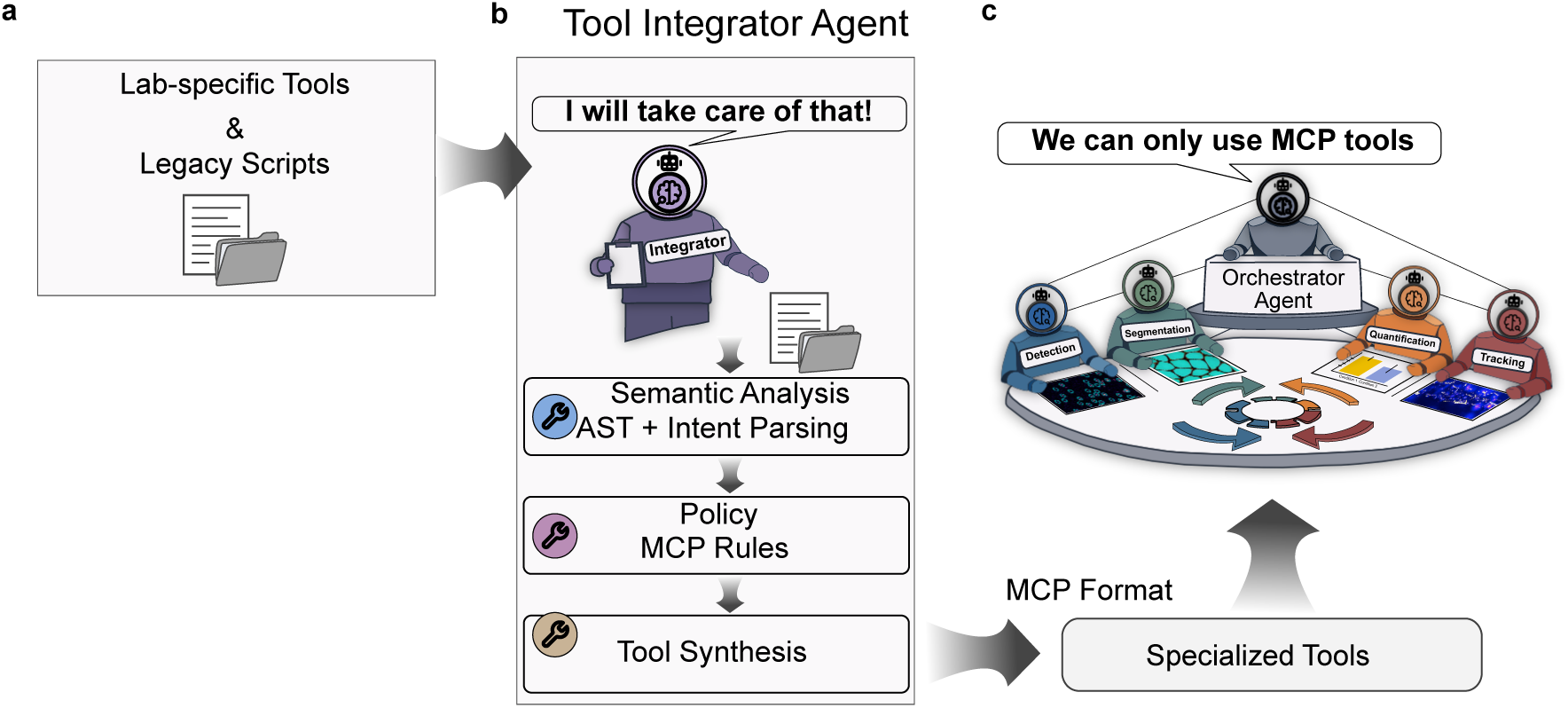
| Automated integration of external analysis scripts into BioBrain-compatible tools. Schematic of the Tool Integrator Agent pipeline. (a) An arbitrary laboratory Python script is provided as input alongside a short functional description. (b) The Tool Integrator Agent performs programmatic code analysis, parsing the script’s structure, identifying public functions, imports, dependencies, and input/output behaviour, and applies a fixed set of integration policies to synthesise a compliant Model Context Protocol (MCP) wrapper with explicit, typed input-output schemas. (c) The resulting tool is registered in the BioBrain MCP tool layer and becomes immediately available to all Sub-Agents through the standard tool-selection mechanism, functionally indistinguishable from manually authored tools.

Once registered, the method becomes indistinguishable from native BioBrain tools: it is accessible to the same Orchestrator agent, subject to the same constrained execution framework, and available through natural-language interaction without further software engineering. This capability enables BioBrain to continuously expand its analytical repertoire as new laboratory methods are developed or published, while preserving transparent and reproducible workflow execution. Rather than requiring experimental scientists to adapt their analyses to existing software, BioBrain adapts existing analytical methods into a common conversational framework.

### Benchmark evaluation across two microscopy modalities

#### Cargo-membrane colocalization analysis

To validate BioBrain on a representative microscopy task we applied it to a simulated two-channel total internal reflection fluorescence (TIRF) dataset modelling cargo release from individual lipid nanoparticles (LNPs). Successful nucleic acid delivery requires cargo release from the nanoparticle, making cargo-membrane dissociation a commonly used proxy for nanoparticle disassembly and release dynamics. In this assay, membrane-labelled LNPs and fluorescent cargo are imaged as diffraction-limited particles in separate channels, and cargo retention is quantified by the fraction of membrane particles that remain spatially colocalized with cargo. The dataset consists of paired single-channel TIFF movies, one cargo channel and one membrane channel, acquired at five discrete timepoints (t0, t30, t60, t90, t120). The simulation models pH-mediated cargo release during endosomal acidification: as pH decreases, LNP structure is destabilised and cargo dissociates from the membrane, producing a monotonic decline in cargo-membrane colocalization over time. This expected monotonic trend provides a useful benchmark as the qualitative biological outcome is known in advance, while the measured colocalization values remain sensitive to realistic image-analysis choices such as particle detection thresholds and association distance. Because the expected biological trend is known a priori while the measured values remain sensitive to image-analysis choices, this dataset provides a controlled benchmark for quantitative evaluation (Table 1).

**Table 1.** | Cargo-membrane colocalization fractions across prompt specificity levels. Values at each timepoint for the simulation ground truth (predefined colocalization), the expert-derived analytical reference, and BioBrain outputs obtained under High-, Mid-, and Low-LOD prompts. The high-level prompt reproduces the expert reference exactly; mid- and low-level prompts show interpretable shifts driven by parameter inference.

| Timepoint | Predefined colocalization | Expert reference | High-LOD prompt | Mid-LOD prompt | Low-LOD prompt |
| --- | --- | --- | --- | --- | --- |
| t0 | 73.68% | 69.6% | 69.6% | 63.3% | 63.8% |
| t30 | 49.97% | 46.8% | 46.8% | 43.2% | 43.5% |
| t60 | 33.95% | 30.9% | 30.9% | 27.8% | 28.1% |
| t90 | 22.96% | 21.3% | 21.3% | 19.1% | 19.2% |
| t120 | 15.55% | 14.3% | 14.3% | 12.2% | 12.2% |

Although the biological objective is straightforward: how does cargo release evolve over time? answering it computationally requires a multi-step workflow. The analyst must handle separate channels, convert data into tool-compatible formats, detect particles independently in each channel, combine of the resulting outputs, and quantify cargo-membrane colocalization across timepoints. These steps often involve multiple tools with different input requirements and data formats, making workflow construction and execution a substantial part of the analytical effort.

Rather than explicitly specifying these operations, the user provided the natural-language request, “*Detect particles in both channels and quantify their spatiotemporal colocalization to assess cargo release*”, From this simple command BioBrain assembled and executed an end-to-end workflow (Fig. 3): The raw two-channel movies were separated into single-channel stacks, each channel was processed independently by a particle-detection tool to localise diffraction-limited objects across frames, and the resulting per-frame coordinates were then passed to a colocalization quantification tool. The colocalization step computed distance-based cargo-membrane association per frame and aggregated results across timepoints to produce a time-resolved release curve and output publication-quality figures.

**Fig. 3.**
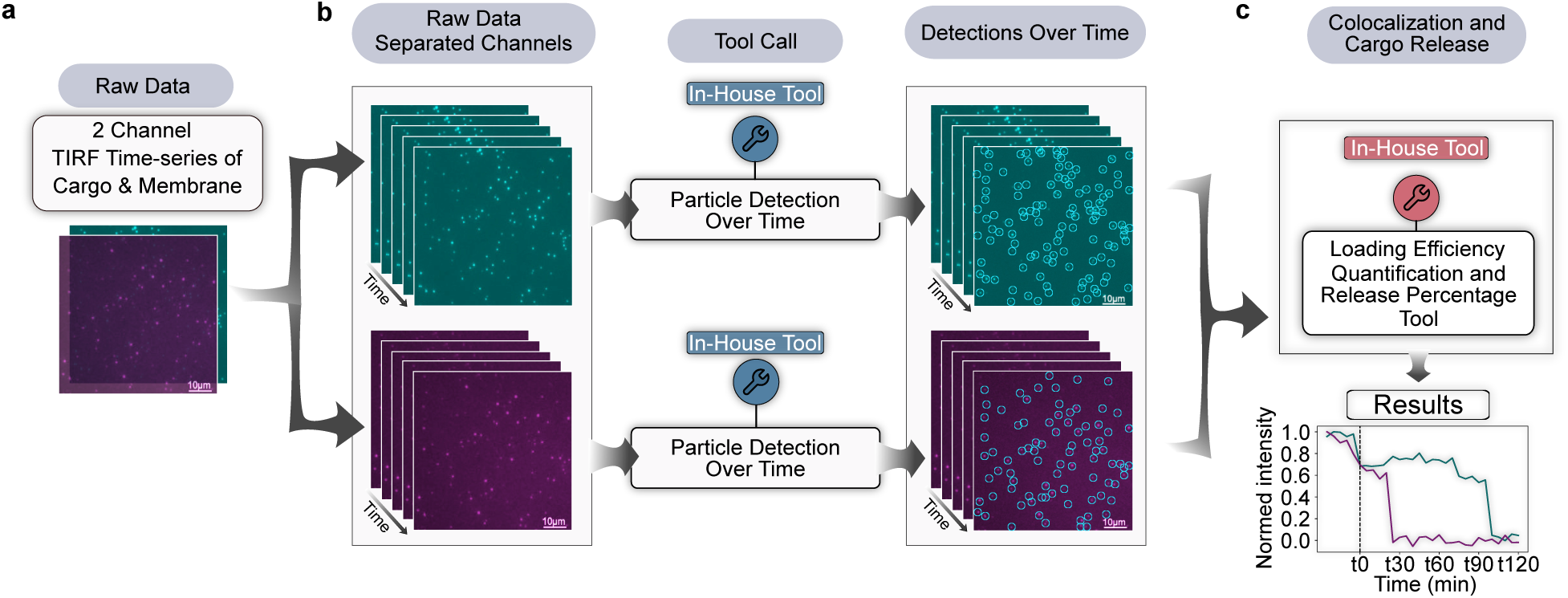
| BioBrain-assembled pipeline for cargo-membrane colocalization analysis in two channel TIRF microscopy. Overview of the BioBrain workflow for quantifying cargo release dynamics from a natural-language instruction. (a) Raw two-channel total internal reflection fluorescence (TIRF) time-series data, displaying Lipid nanoparticles (channel one) loaded with nucleic acid cargo (channel 2). Upon trigger, pH variation cargo is released. Data are provided as input across five timepoints (t0-t120). (b) Each channel is separated and processed independently by the Detection Sub-Agent, which localises diffraction-limited fluorescent particles across imaging frames; representative tool calls and per-frame coordinate outputs are shown. (c) Detected coordinates from both channels are passed to the Colocalization Sub-Agent, which computes pairwise cargo-membrane distances per frame and aggregates results across timepoints, yielding a time-resolved colocalization curve that reflects progressive pH dependent cargo release.

We benchmarked BioBrain against an expert-derived analytical reference, defined as the fully specified workflow a domain data scientist would execute using the same available toolset (see Methods). This reference explicitly specifies every analysis step, including particle-detection settings, cargo-membrane association criteria, and all numerical and parameters values (see Methods). Applied to the simulated dataset, the expert workflow yielded colocalization fractions of 69.6%, 46.8%, 30.9%, 21.3%, and 14.3% from t0 to t120, respectively, closely matching the predefined ground truth. The small offset (1-4%) reflects realistic signal characteristics, including noise, overlapping signals, and imperfect channel separation.

To evaluate robustness to prompt specificity, we repeated the analysis under three levels of user input specificity while holding the dataset and biological objective constant (Supplementary Fig. S4): (i) a fully specified analytical prompt defining tools and numerical parameters explicitly, (ii) a mid-level descriptive prompt conveying intent with qualitative parameter guidance, and (iii) a low-level intent-only prompt of the biological objective (full prompt texts in Methods). Across all prompt levels, BioBrain recovered the correct dataset structure and constructed the same workflow topology: format conversion, particle detection, and distance-based colocalization demonstrating that the underlying computational strategy remained stable despite large differences in prompt specificity.

Under the fully specified prompt, BioBrain reproduced the expert reference outputs verbatim. For the mid-level prompt, BioBrain preserved the correct workflow despite the absence of explicit numerical parameters. It inferred parameter values from the qualitative description, translating phrases such as “small vesicular objects” and “slightly higher SNR” into concrete detection settings (threshold of SNR = 2 and a particle diameter of 9 pixels) while retaining default values for parameters not specified by the user (cargo-membrane association distance of 0.4 pixels rather than the 0.6 pixels specified in the expert reference). These differences produced modest quantitative shifts in measured colocalization but preserved the expected monotonic cargo-release trend. Under the biological-intent-only, low-level prompt, BioBrain identified that essential workflow information was missing, including whether particle detections already existed, and how cargo-membrane colocalization should be operationally defined. BioBrain halted to request clarification rather than proceeding with unsupported assumptions. Once the required information was provided, the workflow was executed using the tool defaults (SNR = 1.0, diameter = 9, distance = 0.4). Despite the minimal analytical guidance, BioBrain recovered the correct workflow structure, preserved the expected temporal release behavior, and maintained full transparency reporting all inferred and default parameter choices, illustrating its fail-safe approach to analytical uncertainty.

Across all prompt levels, BioBrain accurately output the biological trend, differences in quantitative outputs arose from transparent parameter selection rather than changes in workflow construction (Table 1, Supplementary Fig. S5). To assess variability arising from parameter inference, unique to the mid-level, we executed it 100 independent times under identical conditions (Supplementary Fig. S7). Although the resulting colocalization values formed a multimodal distribution, the inferred parameters converged to a small number of discrete combinations rather than varying continuously, with SNR thresholds typically selected from {1.5, 2.0} and particle diameters from {7, 9}. Once selected, each parameter combination produced deterministic and reproducible quantitative outputs, indicating that variability originates from bounded parameter inference rather than stochastic analytical execution.

To benchmark BioBrain’s constrained architecture against direct, unconstrained LLM analysis, we compared it against direct, unconstrained use of two frontier models (Claude Opus 4.8 and GPT-5.5). Each model received the same three prompts, together with a direct path to the dataset and allowed each to generate and execute its own analysis end-to-end (Supplementary Fig. S8).

In contrast to BioBrain, which closely matched the expert-reference values (0-10% deviation) and stayed within 79-95% of the predefined ground truth across all prompt levels, the directly prompted models deviated severely and unpredictably. Counterintuitively, the fully specified high-level prompt produced the largest errors, with both models estimating colocalization levels below 20% of the predefined ground truth, suggesting the models reimplemented the specified detection and colocalization steps in a faithful-looking but quantitatively incorrect manner. Under qualitative prompting the deviations were model-dependent and inconsistent in direction. GPT-5.5, for example, progressively over-estimated colocalization, exceeding the predefined values eventually reversing the expected relationship with the ground truth. Crucially, every run completed without error, producing plausible but incorrect quantitative outputs of exactly the silent-failure type that BioBrain’s constrained architecture is designed to prevent.

Taken together, these results demonstrate that BioBrain reliably recovers the expected cargo-release trend across all prompt conditions, reproduces expert-level quantitative outputs when parameters are fully specified, and degrades predictably when they are not. Variability arises from explicit parameter inference rather than changes in workflow construction or analytical execution, while underspecified requests trigger clarification instead of unsupported assumptions. In contrast, the same prompts issued directly to unconstrained frontier models produced large, model-dependent errors that executed without warning (Supplementary Fig. S8). These findings suggest that the key challenge for AI-assisted quantitative microscopy is not generating code, but translating biological intent into validated, transparent, and reproducible analytical workflows.

### LNP internalization quantification

To test BioBrain on experimental three-dimensional data requiring integration of external tools, we applied it to a two-channel 3D lattice light-sheet microscopy volume containing Jurkat T cells and fluorescently labelled lipid nanoparticles (LNPs). This pipeline exercises capabilities beyond those tested in the TIRF analysis: volumetric rather than temporal processing, true 3D cell segmentation via an external tool, Cellpose, and spatial classification of detected particles relative to segmented cell masks^29,30^. It therefore tests BioBrain’s ability to coordinate heterogeneous tools within a single analysis workflow.

An expert reference workflow segmented cells using Cellpose, detected LNPs using a Crocker-Grier particle detection algorithm, and classified particles as internalized when their centroids fell within the segmented cell masks (see Methods). This analysis identified three cells, 114 LNP particles, 26 internalized particles, and an internalization rate of 22.8%.

From the natural language, intent-level request, *“Segment the cells, detect LNPs, and quantify the number of internalized per cell”*, BioBrain assembled a multi-step volumetric workflow (Fig. 4), involving data parsing, cell segmentation using Cellpose 3D^29^, and LNP-like particle detection using Crocker-Grier and quantifying particle internalization by combining particle coordinates with the segmented cell masks.

**Fig. 4.**
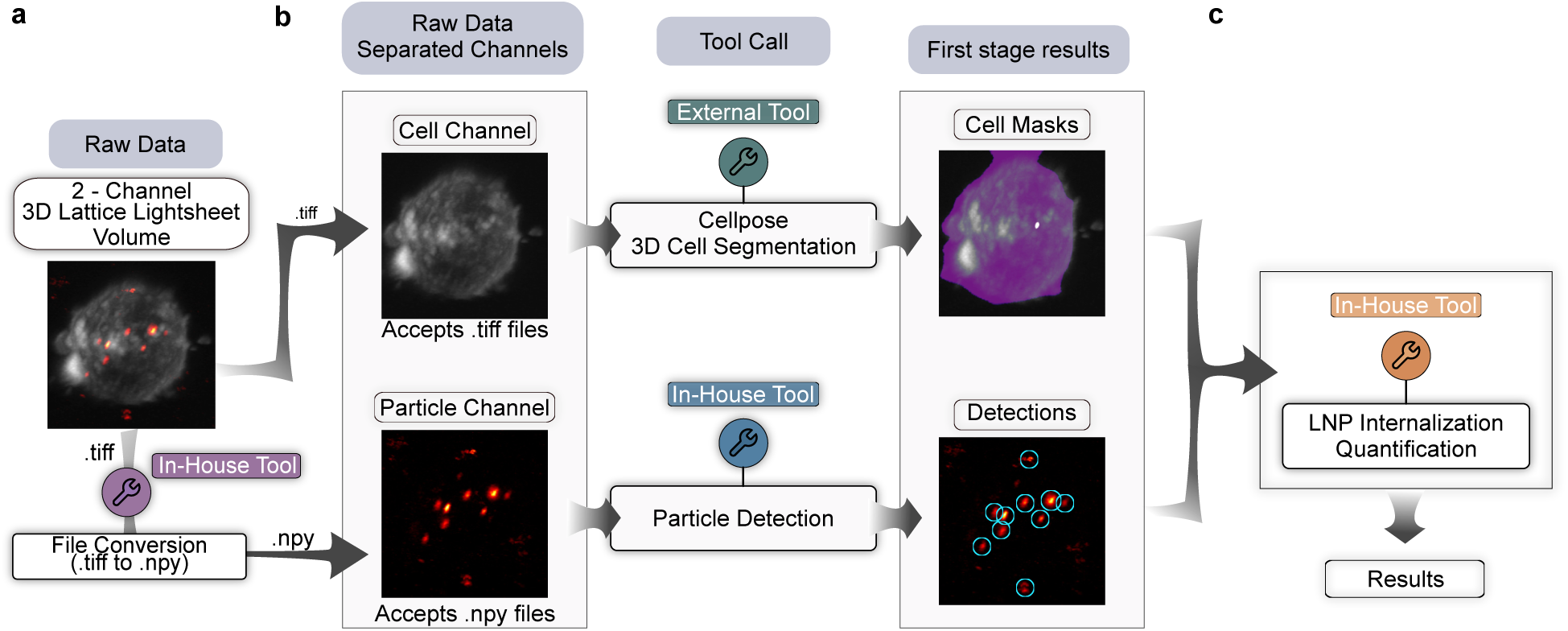
| BioBrain-assembled workflow for quantification of nanoparticles internalization in 3D microscopy data. Overview of the BioBrain pipeline for classifying and counting internalized particles from a natural-language instruction applied to a three-dimensional two-channel microscopy volume. (a) Raw lattice light-sheet data comprising a cell channel and a lipid nanoparticle (LNP) channel are provided as input; BioBrain automatically performs file format conversion to satisfy downstream tool requirements, and a representative volume rendering is shown. (b) The volume is separated into individual channels; the Cell Segmentation Sub-Agent generates three-dimensional cell masks from the cell channel, and the Detection Sub-Agent localises LNP-like objects in three dimensions from the particle channel; first-stage outputs, cell masks and particle coordinate tables, are shown. (c) Cell masks and particle detections are passed to an internalization quantification tool, which classifies each detected particle as internalised or surface-associated based on spatial inclusion within a labelled cell volume and reports per-cell counts and summary statistics.

As with the TIRF pipeline, we evaluated robustness under three prompt specificity levels (Supplementary Fig. S4 and Table S1). Across all three, BioBrain identified the dataset as a single 3D two-channel CZYX volume and constructed the same high-level workflow: format conversion, 3D segmentation, 3D detection, and centroid-in-mask classification. Under the high-level prompt, BioBrain reproduced the expert reference exactly. Under the mid-level prompt, BioBrain preserved workflow structure and tool choices but inferred the detection threshold from qualitative guidance, “slightly higher SNR than default”, yielding 181 detected particles, 22 internalized, 18.2% internalization rate. Under the low-level prompt, BioBrain correctly identified that the channel identities of cells and nanoparticles were not specified, and halted to request clarification rather than guessing. After clarification, BioBrain proceeded with default detection settings, yielding 1,278 detected particles, 167 internalised, and a 13.1% internalization rate (See Table 2).

**Table 2.**
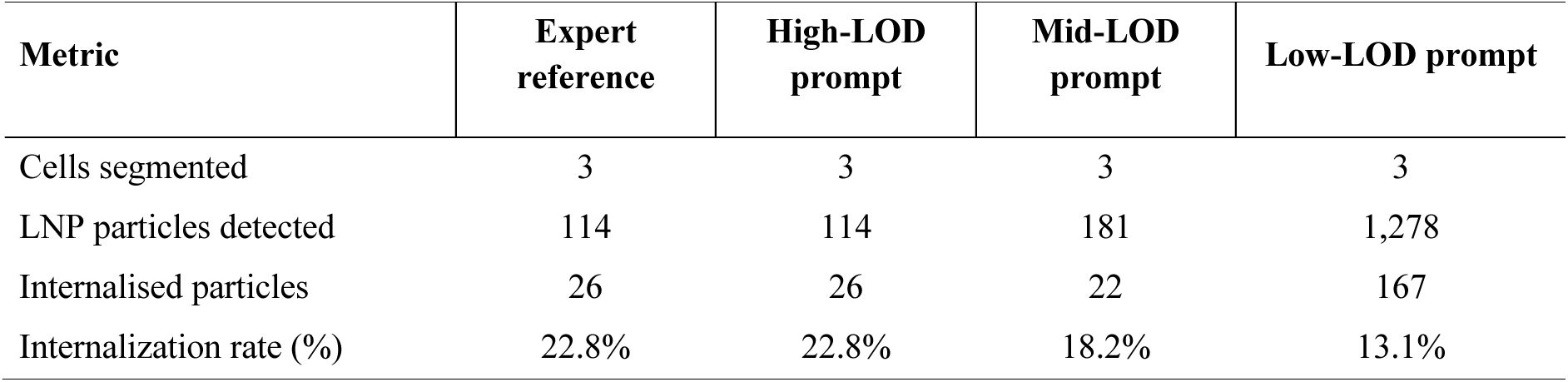
| LNP internalization metrics across prompt specificity levels. Number of segmented cells, detected LNP particles, internalised particles, and overall internalization rate for the expert reference and BioBrain outputs under High-, Mid-, and Low-LOD prompts.

Importantly, cell segmentation remained stable across all prompt conditions (Supplementary Fig. S6), whereas differences in quantitative outputs originated almost entirely from particle detection parameterization. Lower detection thresholds admitted a substantially larger population of low-intensity particles distributed throughout the imaging volume, increasing total particle counts more strongly than internalized counts and thereby reducing the calculated internalization ratio. Because BioBrain reports intermediate analytical outputs together with final summary metrics, the source of this deviation could be traced directly to particle detection rather than segmentation or particle-to-cell assignment.

Together, these two benchmarks span complementary regimes, temporal two-dimensional data and volumetric three-dimensional data, demonstrate that BioBrain is not tied to a particular task type but generalizes across complementary microscopy modalities. In essence, BioBrain holds the potential to operate for any analysis for which suitable tools exist. Across both applications, expert-derived results were reproduced when analytical parameters were specified, whereas deviations under underspecified prompts arose from transparent parameter choices rather than workflow construction errors. These findings suggest that BioBrain’s reliability stems from constraining AI to validated analytical workflows while exposing every computational decision to the user. It also demonstrates that AI can reliably translate biological intent into quantitative microscopy analysis by constraining reasoning to validated computational workflows.

## Discussion and conclusion

Current advances in microscopy have radically expanded the range of biological questions that can be addressed, yet the ability to extract quantitative information from these data has not advanced at the same pace and across fields. Increasingly, the limiting factor is not image acquisition but the expertise required to construct and execute complex computational workflows. BioBrain addresses this asymmetry by providing a natural-language interface that automatically, rapidly and precisely translates biological intent into validated and reproducible image-analysis pipelines. Our benchmarks across two complementary microscopy modalities, demonstrate that the BioBrain framework reproduced expert-derived analyses when parameters were specified and produced transparent, traceable deviations when user input was incomplete, requesting clarification rather than proceeding under unsupported assumptions.

Our results and benchmarking indicate that the reliability of AI-assisted scientific analysis depends less on the capabilities of the underlying language model than on the architecture within which it operates. Direct use of frontier language models generated plausible but quantitatively incorrect analyses without explicit failure^27^, whereas BioBrain constrained reasoning to validated analytical methods and deterministic tool execution. Consequently, variability was confined to explicit parameter inference rather than workflow construction, and every analytical decision remained inspectable and reproducible. More broadly, these findings indicate that reliable scientific AI, today, requires translating biological intent into validated computational workflows rather than relying on unconstrained code generation. Importantly, this architecture keeps raw data and tool execution local: the language model receives the user prompt and tool descriptions, but not the microscopy data, intermediate outputs or tool source code. Because the framework is model agnostic, the same workflow can also be run with a local language model when offline analysis is required.

Besides enabling natural-language analysis, BioBrain addresses a broader challenge facing the bioimaging community: the growing disconnect between published computational methods and their practical accessibility. Although many state-of-the-art algorithms are openly released, they are often embedded in laboratory-specific scripts and software environments^19^ that are difficult for non-specialists to install, adapt, and integrate into new analytical workflows. Consequently, valuable analytical advances often remain confined to the laboratories that developed them. By automatically converting existing scripts into standardized analytical tools, BioBrain provides a mechanism for incorporating both legacy and newly developed methods into a common conversational framework. In this way, analytical innovations can become immediately accessible to experimental scientists without requiring them to reconstruct complex computational environments, potentially advancing the FAIR principles of research software^31^. More broadly, BioBrain acts as a translation layer between the rapidly expanding ecosystem of computational microscopy methods and the experimental scientists who seek to apply them, helping to bridge the gap between methodological innovation and biological discovery.

Several boundaries of the current framework follow directly from its design rather than from incidental gaps. Because BioBrain executes only validated tools rather than generating arbitrary analytical code, its capabilities is set by the methods available in its toolset. This constraint is also the source of BioBrains reliability: workflow construction is limited to established analytical methods, preventing unsupported computational steps. Coverage, not reliability, is therefore the axis along which the system grows: the Tool Integrator Agent provides a direct path to extend BioBrain to particle tracking, morphological profiling, spectral imaging, and further future developed domains as validated methods become available, without altering its reliability and reproducibility. Similarly, variability arising from qualitative prompts is an inherent consequence of translating the biological intent into quantitative parameters. this variability is confined to a transparent parameter-inference step, while the analytical workflow and tool execution remain deterministic. Every inferred parameter is reported alongside the results, allowing users to distinguish between specified and inferred choices and to increase prompt specificity when quantitative precision is required. Importantly, this variability is limited to the choice of analysis parameters, while the underlying analytical workflow and computational methods remain unchanged.. Continued benchmarking across language models and tasks, together with laboratory-specific memory, standardized provenance tracking, and integration with community resources such as the BioImage Model Zoo, would further extend reproducibility and reach^22,23,32^.

Ultimately, the significance of BioBrain lies in changing how scientists interact with existing image-analysis methods, rather than replacing them. As imaging technologies continue to grow in complexity and throughput, the gap between data acquisition and biological interpretation will widen unless analytical tools become more accessible. BioBrain demonstrates that LLM-driven reasoning can be combined with constrained, deterministic execution to enable experimental scientists to communicate with computational tools in the language of biology rather than the language of software, providing a practical path toward democratizing quantitative microscopy and accelerating biological discovery.

## Methods

### BioBrain system implementation

BioBrain is implemented in Python and built on the Google Agent Development Kit (ADK), which provides the framework for defining, initialising, and coordinating all agents. The orchestrator agent and all domain-specific sub-agents are powered by the DeepSeek V3 chat model (non-reasoning variant), selected for its balance of capability and cost-efficiency. Importantly, the framework is model-agnostic by design: different LLMs can be assigned to different agents depending on task requirements. For example, the Tool Integrator Agent, which must parse and wrap arbitrary code, could alternatively employ a model optimised for code understanding. All LLM calls use a sampling temperature of 0.2 to promote deterministic, reproducible agent behaviour across runs. Inter-agent data flow is coordinated by the orchestrator through the declared input-output requirements of each tool: the output schema of one step must satisfy the input schema of the next, ensuring type-safe composition of multi-step workflows.

Analysis tools are implemented as deterministic Python functions with typed arguments and descriptive docstrings. Tool schemas are defined using Python function signatures and type hints, which are automatically converted into JSON Schema when tools are exposed through Model Context Protocol (MCP) servers. MCP servers are implemented using the FastMCP Python package with standard I/O (stdio) transport, chosen for its low-latency performance in single-system deployments where the entire BioBrain repository resides on one machine. Agent memory is implemented as a simple conversation history maintained within each session; cross-session memory is not currently retained. All experiments reported in this study were performed on a MacBook Pro equipped with an Apple M3 Pro chip, except for the lattice light-sheet internalization pipeline, which required GPU acceleration for three-dimensional cell segmentation and was executed on a workstation equipped with an NVIDIA RTX 5000 GPU. Typical end-to-end execution times were approximately 6-7 minutes for the cargo-membrane colocalization pipeline and 8-9 minutes for the LNP internalization pipeline (on the GPU workstation). BioBrain is distributed as a containerised software environment in which the full local codebase, including the orchestrator, sub-agents, MCP servers, tools, repositories, and dependencies, is built into a single Docker image. This design provides a reproducible execution environment for the complete multi-agent system. Users interact with BioBrain through the Google ADK web interface, launched locally from the terminal, where natural-language prompts are submitted and the returned results and analytical decisions can be reviewed. The code will be made available upon publication. A summary of all system configuration parameters is provided in Supplementary Table S2.

### Tool Integrator agent

The Tool Integrator Agent automates the conversion of arbitrary Python analysis scripts into MCP-compliant tools, removing the need for manual schema authoring or data-science intervention during tool onboarding. When a new script is placed in a designated input directory, the agent inspects it using the Python ast (Abstract Syntax Tree) standard-library module to perform programmatic code analysis. This inspection identifies public functions, import statements, external dependencies, file input/output behaviour, and candidate entry points. On the basis of this analysis, the agent generates a standardised MCP wrapper that exposes exactly one public function per tool, enforces strict input-output schemas with explicit type annotations, follows deterministic import and dependency resolution rules, and registers the resulting tool directly in the BioBrain MCP tool server. External dependencies required by the integrated script are automatically installed at integration time via a dedicated dependency-installation tool.

Each generated wrapper adheres to the same schema and dependency conventions as manually authored tools, ensuring that automatically integrated tools are indistinguishable in interface and reliability from hand-crafted components. The wrapper synthesis is governed by a fixed set of integration policies that constrain the output format, prohibit dynamic code generation within the wrapper, and require that all tool parameters be explicitly typed and documented.

After wrapper generation, BioBrain performs an automated validation step before registering the tool in the system. First, the generated wrapper is checked for interface correctness, ensuring that the tool exposes a valid schema derived from the function signature (argument names, types, and required fields). Next, the tool is executed on synthetic or minimal test inputs generated by the Tool Integrator Agent to verify that it runs without runtime errors and that the returned output conforms to the expected structure. During this step, missing dependencies are automatically installed if required. Once validation passes, the new tool is immediately available as an MCP tool trough the tool integrator agent.

The Tool Integrator Agent currently supports single-file Python scripts with a clearly identifiable entry point. Scripts comprising multi-file packages or containing multiple independent functionalities without a single clear entry point are not automatically handled; in such cases, BioBrain prompts the user to specify which function should be exposed. Scripts with graphical user interface dependencies or global mutable state are also outside the current scope of automatic integration.

### Simulated TIRF colocalization dataset

The simulated two-channel TIRF time-series dataset was generated using custom Python code (NumPy for array operations, tifffile for saving TIFF stacks). No external simulation framework was used. Each movie consists of 20 frames at 512 × 512 pixels. Five timepoints (t0, t30, t60, t90, t120) represent minutes and model endosomal acidification during receptor-mediated endocytosis: as endosomes mature and luminal pH drops, cargo dissociates from membrane receptors, reducing colocalization according to a geometric decay.

Particles are placed at uniformly random integer pixel positions (x, y drawn independently). The membrane channel contains 100-300 particles per frame (uniform random draw), and the cargo channel contains 200-400 particles per frame. Each particle is rendered as a two-dimensional isotropic Gaussian with σ drawn uniformly from 0.3-1.0 pixels (corresponding to a full-width at half-maximum of approximately 0.7-2.4 pixels) and peak amplitude drawn uniformly from 300-400 intensity units. No Airy disk model is used.

Colocalization at each timepoint is enforced by exact pixel coincidence: a subset of membrane-channel coordinate tuples is copied directly into the cargo-channel coordinate list, so paired particles occupy identical pixel positions. The predefined cargo-membrane association rates are 74.0% (t0), 50.3% (t30), 34.0% (t60), 23.0% (t90), and 15.6% (t120).

A mixed Poisson-Gaussian noise model is applied. First, Poisson shot noise is applied to the signal image, then additive Gaussian readout noise with a standard deviation of 5 intensity units is added. Given the particle amplitude range of 300-400 and the Gaussian noise scale, the approximate peak signal-to-noise ratio is on the order of 60-80 before Poisson noise.

### Sort LNP formulation

SORT LNPs were formulated as previously described^33^. Lipids were mixed at molar ratios of 50/10/38.3/1.5/0.2 SM-102/DSPC/cholesterol/DMG-PEG2000/ATTO655-DOPE, with DOPS added at 10% weight/weight ratio relative to total lipid (Avanti Polar Lipids and ATTO-TEC). All lipids were dissolved in 99% ethanol with the exception of DOPS, which was reconstituted in THF. mRNA cargo (CleanCap EGFP mRNA, fully 5-methoxyuridine-substituted; TriLink BioTechnologies) was solubilized in 10 mM RNase-free citrate buffer at pH 4.0 immediately before LNP assembly. LNPs were assembled using a microfluidic device (Harvard Apparatus Pump 33 DDS) fitted with a herringbone mixer glass chip at a 1:40 weight/weight ratio of mRNA:lipid, a 3:1 aqueous-to-ethanol flow-rate ratio, and a total flow rate of 500 µL min⁻¹. Ethanol was removed by diluting the sample in cold PBS (1×) to a total volume of 4 mL in a 10 kDa molecular-weight-cutoff Amicon Ultra centrifugal filter (MilliporeSigma) and diafiltering by centrifugation at 3,000 × g at 4 °C. Dilution and diafiltration were performed twice, after which samples were stored at 4 °C. The physical properties of the SORT LNPs, specifically size and polydispersity, were assessed via dynamic light scattering using a Malvern Panalytical Zetasizer; a red filter was employed to prevent interference from ATTO655 fluorescence. To quantify mRNA loading and total concentration, a Quant-it™ RiboGreen RNA Assay Kit (Invitrogen) was utilized. Following the manufacturer’s low-range guidelines, unencapsulated mRNA was measured in intact particles and compared against total mRNA released was measured after disruption of LNPs with 0.1% v/v Triton X-100. All fluorescence measurements were captured on a SpectraMax iD3 plate reader, and experimental dosing was standardized based on the total encapsulated mRNA content.

### Cell culture and plating

Jurkat cells (Cytion/CLS, ref 302147) were utilized for all experiments. Cells were cultured in RPMI1640 (Sigma-Aldrich) with 10% heat-inactivated foetal bovine serum (FBS, Sigma-Aldrich) at 37°C and 5% CO2. Cells were used between split levels 6-21.

### Cell Seeding and Experimental Setup

On the day of treatment, Jurkat T cells were seeded at a density of 20,000 cells per well into 8-well glass-bottom-slides (Ibidi). To facilitate cell attachment, the slides were pre-coated with human fibronectin (Sigma-Aldrich) in DPBS for 1 hour at room temperature, followed by three DPBS washes. Prior to LNP addition, cells were suspended in serum-free RPMI 1640 (Fisher Scientific) and allowed to equilibrate for 60 minutes.

### Lattice light-sheet microscopy

Imaging was performed on a Lattice Lightsheet 7 microscope (ZEISS) equipped with two ORCA-FusionBT sCMOS cameras (Hamamatsu). Data were acquired using the 30 × 1,000 µm light sheet. Before each imaging session, the light sheet was aligned by casting 0.1 µm TetraSpeck fluorescent microspheres (Thermo Fisher Scientific) in high-gel-strength agarose (1:100 dilution) in an empty well and optimising the point spread function. The isotropic voxel size after deskewing was 0.145 × 0.145 × 0.145 µm.

Cells were stained with CellMask Green plasma-membrane stain (1:1,000 dilution; Sigma-Aldrich) for 5 min, washed once in serum-free RPMI 1640 (Fisher Scientific), and then treated with SORT LNPs. CellMask Green was excited with a 488 nm laser at 1% power with 5.0 ms exposure; ATTO655 (LNP label) was excited with a 640 nm laser at 3% power with 5.0 ms exposure. Volumes were acquired at 200 nm z-step intervals over 315 slices per field of view for 25 time frames. Imaging was performed at 37 °C with 5% CO₂ atmosphere control. A cover-glass transformation (ZEISS) was applied to the raw data before analysis. A single representative volume (first time frame) was used for the BioBrain internalization analysis.

### Expert reference workflows

To benchmark BioBrain’s outputs, we defined expert-derived analytical reference workflows for each pipeline, representing the analysis a domain data scientist would execute using the same toolset available to BioBrain. These references specify every processing step, tool, and parameter value, and serve as the primary comparison for all evaluations.

#### Cargo-membrane colocalization

Starting from paired single-channel TIFF movies at each of five timepoints (t0, t30, t60, t90, t120), the expert reference workflow proceeds as follows. Both cargo and membrane movies are first converted from TIFF to NumPy array format to satisfy the input requirements of the particle-detection tool. Particle detection is then performed independently on each channel using a Crocker-Grier-based detector with a particle diameter of 7 pixels and a signal-to-noise ratio (SNR) threshold of 1.2, treating each movie as a 2D time series^34,35^.. Colocalization is quantified per timepoint by computing pairwise Euclidean distances between cargo and membrane detections and labelling a membrane-channel particle as colocalized if a cargo-channel particle is found within a maximum distance of 0.6 pixels. This threshold was selected to accommodate sub-pixel registration offsets between channels while remaining below the diffraction-limited spot diameter, ensuring that only spatially coincident particles are classified as colocalized. The output consists of per-particle colocalization annotations and per-timepoint summary statistics (number and percentage of colocalized particles). Applied to the simulated dataset, this reference yields colocalization fractions of 69.6%, 46.8%, 30.9%, 21.3%, and 14.3% at t0 through t120, respectively.

#### LNP internalization

The input is a single two-channel three-dimensional OME-TIFF volume with axis order (C, Z, Y, X) and no time dimension, where channel 0 contains LNP signal and channel 1 contains membrane-stained Jurkat cells. Cells are segmented in three dimensions by applying Cellpose (version 4.0.4, default model in 3D mode) to channel 1 with a minimum cell size of 25 voxels and a minimum cell diameter of 20 voxels, producing a labelled three-dimensional mask in which each cell receives a unique integer identifier^29,30^. In parallel, the volume is converted to NumPy format to satisfy the detection tool’s input requirements. LNPs are detected on channel 0 using a Crocker-Grier-based method with an SNR threshold of 1.7 and a particle diameter of 7 voxels, yielding a table of (z, y, x) centroids and per-particle intensity measurements^34^. Internalization is quantified by classifying a detected particle as internalised if its centroid, rounded to the nearest voxel (no sub-voxel interpolation), falls within a labelled voxel of the cell mask. The output comprises per-particle internalization annotations, per-cell internalised counts, and global summary statistics. Applied to the dataset used in this study, the expert reference yields 3 segmented cells, 114 detected LNP particles, 26 internalised particles, and an internalization rate of 22.8%.

### Prompt specifications for benchmark evaluation

We evaluated BioBrain on each pipeline using three levels of prompt specificity, holding the dataset and analytical objective constant while systematically varying the amount of technical detail provided. The three levels are defined as follows.

#### High-level analytical prompt

The user specifies the exact tools, algorithms, and numerical parameters to be used at each step (e.g., particle diameter, SNR threshold, association distance). This level tests whether BioBrain can faithfully execute an explicitly defined workflow.

#### Mid-level descriptive prompt

The user describes the analytical intent with qualitative parameter guidance (e.g., “small vesicular objects”, “slightly higher SNR than default”) but omits specific numerical values and algorithm names where possible. This level tests BioBrain’s ability to infer reasonable parameters from domain-relevant language.

#### Low-level intent-only prompt

The user states only the analytical objective with minimal technical detail (e.g., “What is the percentage of colocalized particles per timepoint?”). This level tests BioBrain’s ability to autonomously identify the required processing steps and request clarification when essential information is missing.

The complete prompt texts for both pipelines at all three levels are provided in Supplementary Table S1. For each prompt level, we compared BioBrain’s outputs against the expert reference on the following dimensions: workflow topology (whether the same sequence of processing steps was constructed), tool selection (whether the same tools were chosen), parameter values (exact match, inferred value, or tool default), clarification behaviour (whether and when BioBrain halted to request user input), and quantitative outputs (colocalization fractions or internalization rates).

### Variability analysis under qualitative prompting

Because the mid-level prompt requires the LLM to infer numerical parameters from qualitative descriptions, this condition introduces a source of run-to-run variability absent in the fully specified (high-level) and default-driven (low-level) conditions. To characterise this variability, we executed the mid-level prompt for the cargo-membrane colocalization pipeline 100 independent times under identical conditions (same LLM model version, API settings, temperature, dataset, and file paths) and recorded the resulting colocalization percentages at each timepoint. All 100 runs were performed sequentially; no post-hoc filtering was applied (no runs were excluded for failure or timeout).

The variability analysis was performed only for the TIRF colocalization pipeline because the primary objective was to quantify the variability in LLM-inferred detection parameters (SNR threshold and particle diameter) under qualitative prompting. Since the colocalization pipeline isolated this source of variability, repeating the analysis for the LNP internalization pipeline was unnecessary: the detection parameter space is the same, and the additional ambiguity introduced by cell segmentation parameters was not expected to produce meaningful variability given the clearly defined cell morphology in the lattice light-sheet data, where modest parameter perturbations yield identical segmentation masks.

#### Direct LLM baseline

To compare BioBrain’s constrained architecture against direct, unconstrained large language model analysis, we evaluated two frontier models, Claude Opus 4.8 (Anthropic) and GPT-5.5 (OpenAI), on the cargo-membrane colocalization task. Each model was given the identical three prompts used for the BioBrain benchmark (High-, Mid-, and Low-LOD; full texts in Supplementary Table S1), together with a direct filesystem path to the same simulated TIRF dataset. Unlike the BioBrain runs, no curated tools, schemas, or orchestration layer were provided: each model was operated in an agentic code-execution mode using Claude code or codex with filesystem and Python execution access, in which it could read the data, and autonomously write and execute its own analysis code to produce per-timepoint colocalization percentages. Models were queried with default sampling temperature; no analytical guidance beyond the benchmark prompt was given, and no intermediate outputs were corrected.

Each model was run 1 time per prompt level, and the resulting per-timepoint colocalization percentages were printed by the executed code. Outputs were compared against the predefined colocalization rates and the expert reference on the same metrics used for BioBrain (per-timepoint colocalization fraction and the monotonic-decline trend).

## Author contributions

A.B., K.T. and N.S.H. wrote the manuscript with feedback from all authors. K.T. and S.M. prepared the figures with the help of A.B. and N.S.H.. K.T. conceived and initiated the project, designed the multi-agent framework architecture, and developed all agents and analysis tools with feedback from NSH. A.B. developed the colocalization analysis tools. S.B., M.W.D., and A.B. provided critical input on system design and analytical workflows throughout development. A.O. contributed to conceptual development and co-developed the three-dimensional cell-mask segmentation pipeline with K.T.. K.T. performed all benchmark experiments and analysed the data with help from A.B., S.M. and N.S.H.. N.S.H. had the overall supervision and project management, and acquired funding.

## Acknowledgements

We thank Sara Vogt Bleshøy for providing the Jurkat T cell and lipid nanoparticle lattice light-sheet microscopy data used in this study.

## Disclaimer

N.S.H. is the CSO and co-founder of EDGE Biotechnologies. K.T. and A.B. are part-time employees of EDGE Biotechnologies.

