## supplementary information for "BioBrain: A Multi-Agent Framework for Natural Language Driven Quantitative Microscopy Data Analysis"

### Supplementary Figures

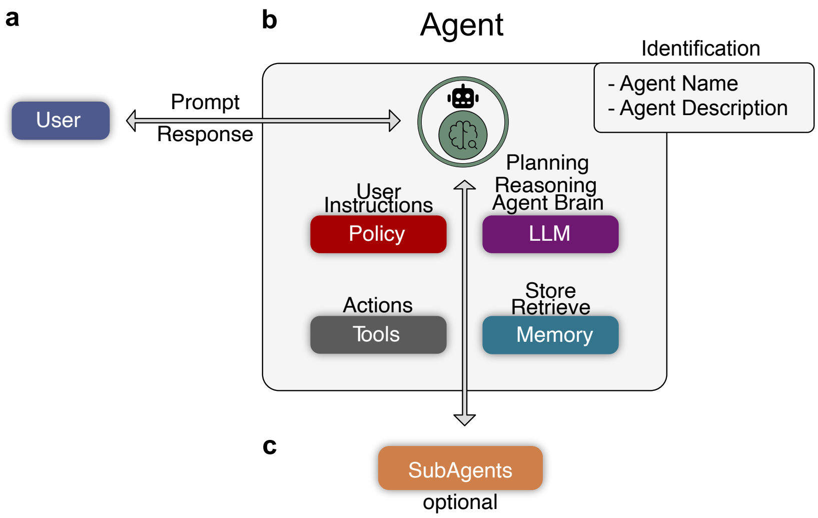


**Supplementary Fig. S1 | Core agent architecture of the BioBrain framework.** Schematic representation of the fundamental components of a BioBrain agent. (a) The user communicates with the BioBrain system through bidirectional natural-language exchange, submitting analytical prompts and receiving structured results and explanations in return. (b) Each agent in BioBrain shares a common internal architecture: a dedicated name and functional description enable programmatic recognition and selection by other agents; a policy module encodes predefined instructions and operational constraints; a large language model (LLM) serves as the reasoning core, performing planning and decision-making; a tool interface translates high-level reasoning into concrete computational operations; and a memory module stores contextual information, including previous actions and intermediate results, to support multi-step, stateful workflows. (c) When required, any agent can delegate subtasks to specialised sub-agents, forming a hierarchical structure in which reasoning and execution responsibilities are distributed across the system.

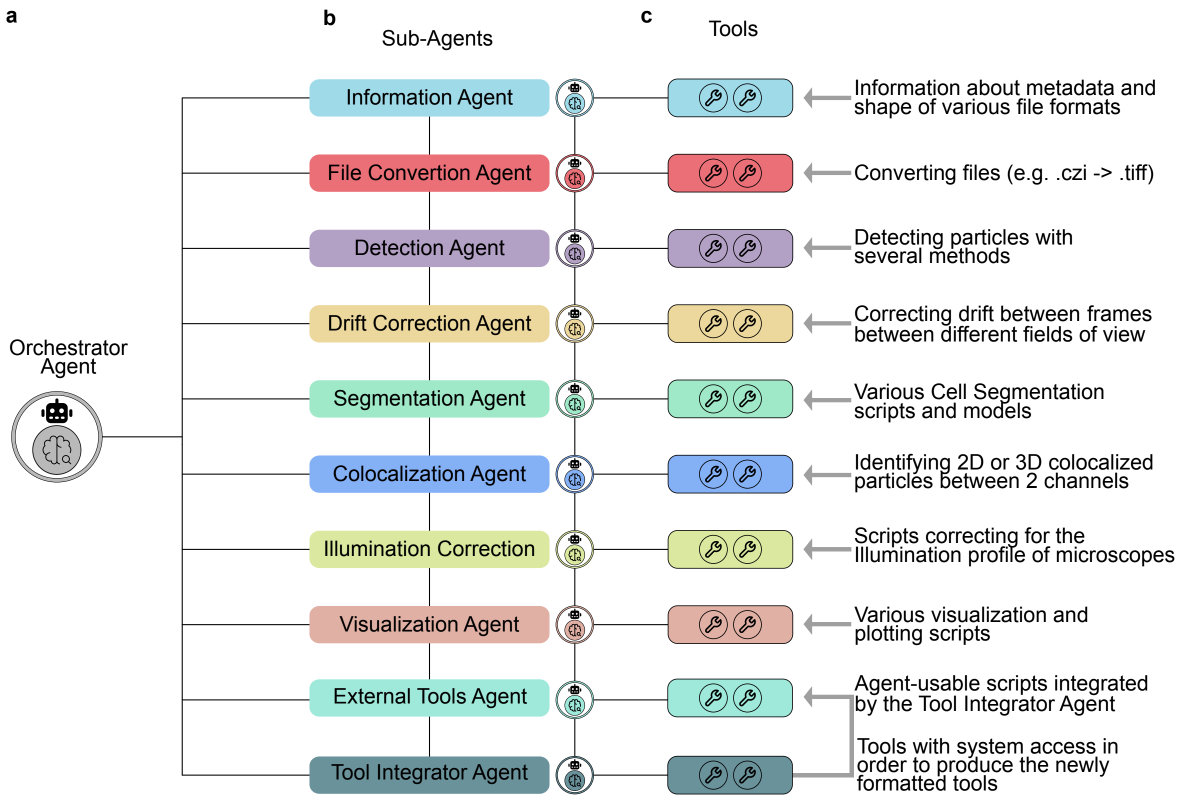


**Supplementary Fig. S2 | Hierarchical multi-agent architecture of the BioBrain framework.** Schematic overview of the full BioBrain system architecture. (a) The Orchestrator Agent receives natural-language instructions from the user and dynamically routes subtasks to the appropriate Sub-Agent based on each agent's name and functional description. (b) The full set of domain-specific Sub-Agents available within BioBrain, each operating within its own analytical scope. (c) Representative toolsets associated with each Sub-Agent, comprising both in-house and external tools grouped by functional category rather than by individual tool name. Tools within each category may include multiple specific implementations; for a full description of available tools and their specifications, see Supplementary Table S3.

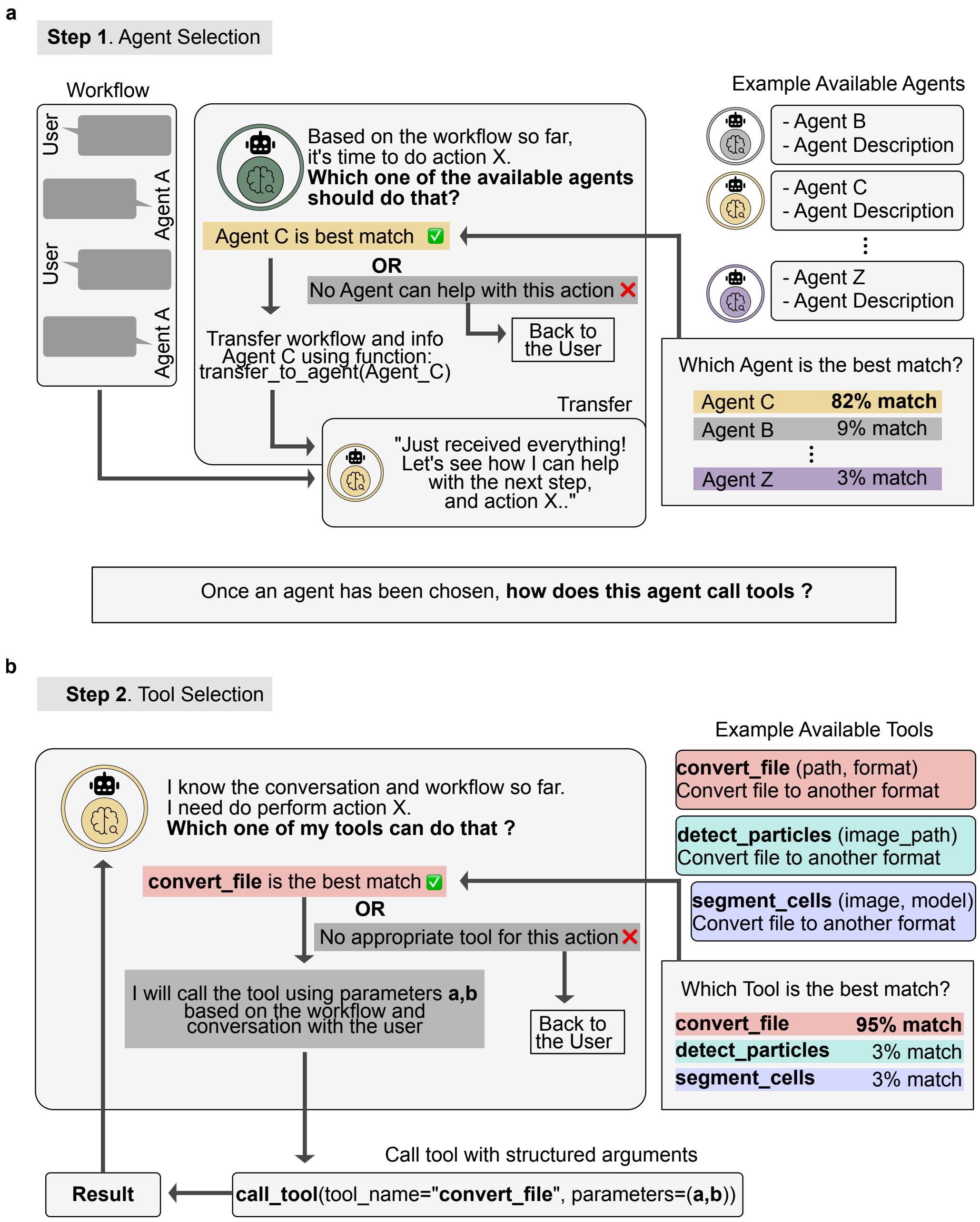


**Supplementary Fig. S3 | Two-step decision loop governing agent delegation and tool selection in BioBrain.** Schematic of the runtime process by which BioBrain resolves a user request into delegated agent responsibilities and parameterised tool calls. **(a)** Agent selection: the active agent evaluates the next required action against the catalogue of available agents, each defined by a name and functional description, and delegates to the best-fit candidate, transferring full workflow context to ensure continuity. If no suitable agent is found, control returns to the user for clarification. **(b)** Tool selection: the delegated agent applies the same matching logic within its own toolset, ranks candidate tools against the requested action and current context, determines the required parameters, and invokes the selected tool through a structured call. The tool returns a result that is propagated back to the agent for downstream use. At both steps, if no sufficient match is found, BioBrain halts and requests clarification rather than proceeding under uncertainty.

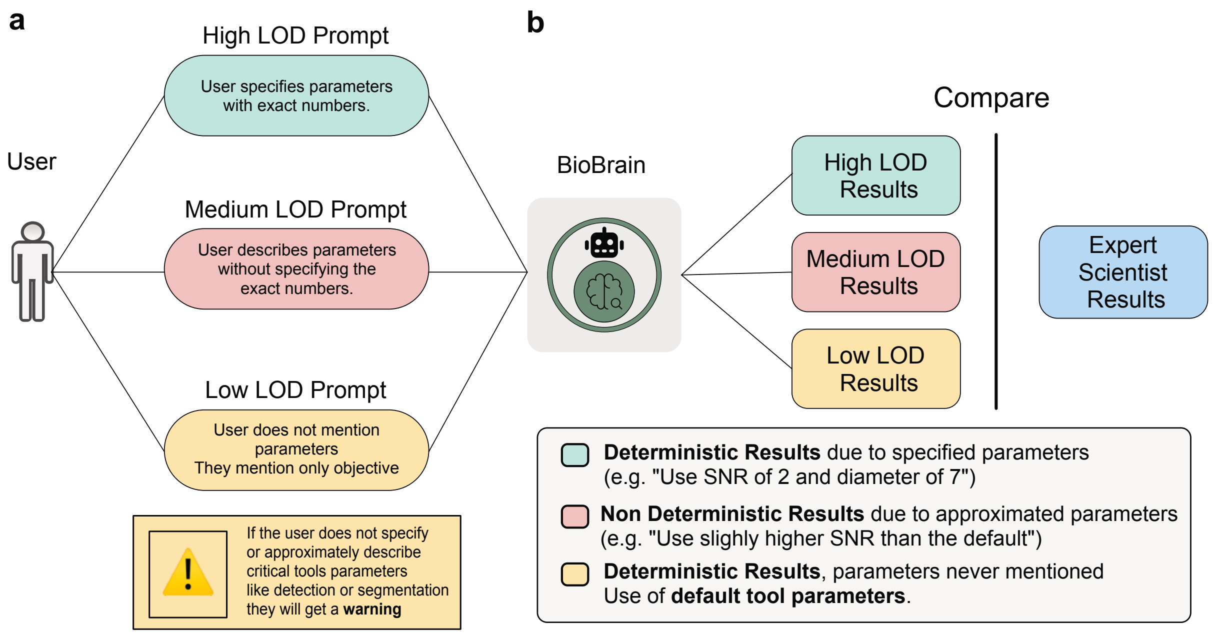


**Supplementary Fig. S4 | Evaluation framework for assessing the effect of prompt specificity on BioBrain outputs.** Schematic of the experimental design used to evaluate how prompt specificity influences BioBrain's parameter selection and analytical outputs. **(a)** Three prompt specificity levels evaluated in parallel: High Level of Detail (High-LOD), in which exact tools and numerical parameter values are specified; Medium Level of Detail (Mid-LOD), in which analytical intent is described qualitatively; and Low Level of Detail (Low-LOD), in which only the analytical objective is stated. **(b)** BioBrain outputs at each prompt level are compared against an expert-derived analytical reference.

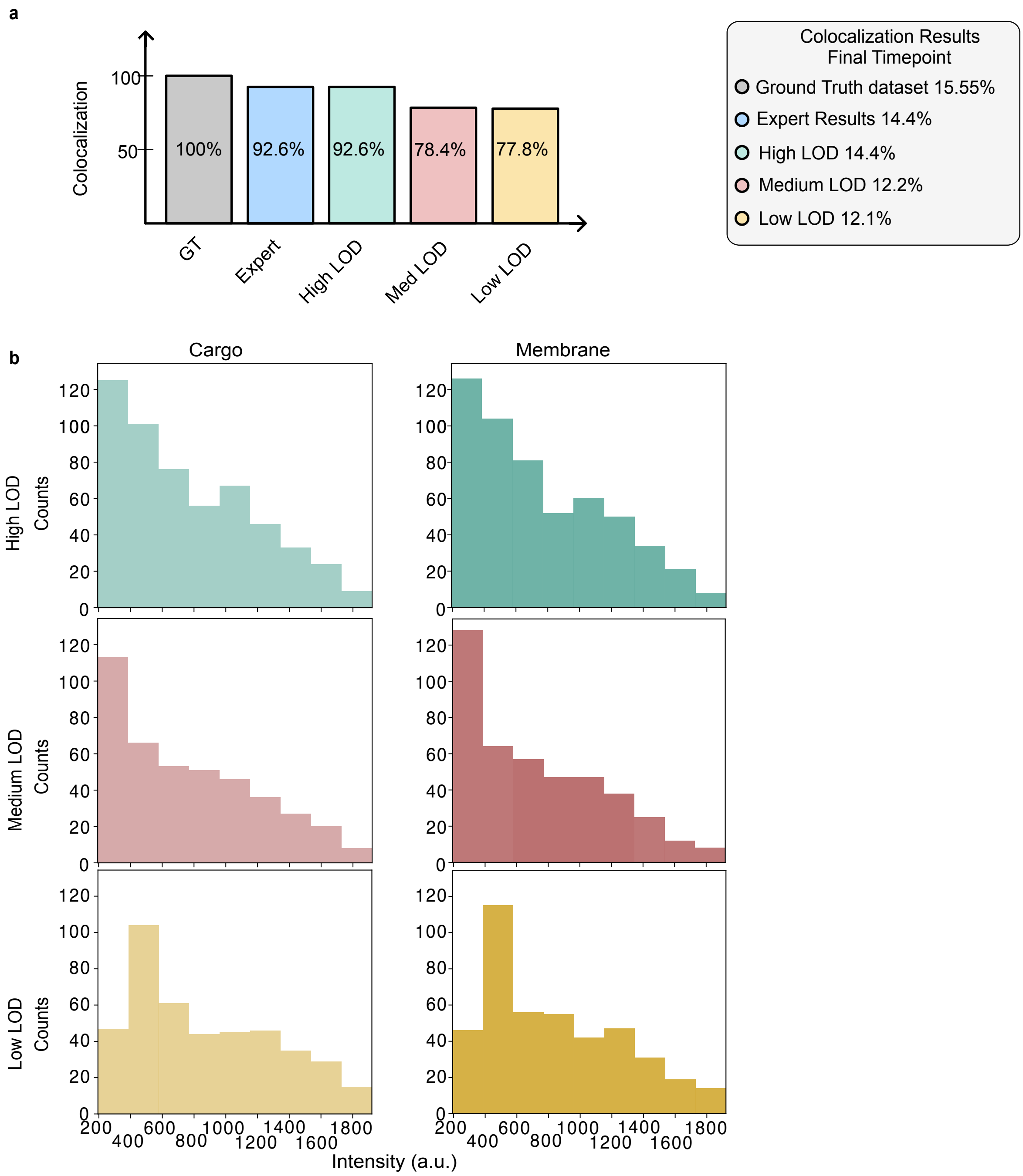


**Supplementary Fig. S5 | Cargo–membrane colocalization outputs across prompt specificity levels.** (a) Bar plot of colocalization fraction at the final timepoint (t120) for the simulation ground truth (GT), the expert analytical reference, and BioBrain outputs obtained under High-, Mid-, and Low-LOD prompts. Values are normalised to the predefined ground-truth colocalization rate; absolute colocalization fractions for each condition are reported in the legend**.** (b) Intensity distributions of particles classified as colocalized, shown for the cargo channel (left column) and membrane channel (right column) under High-, Mid-, and Low-LOD prompts (rows).

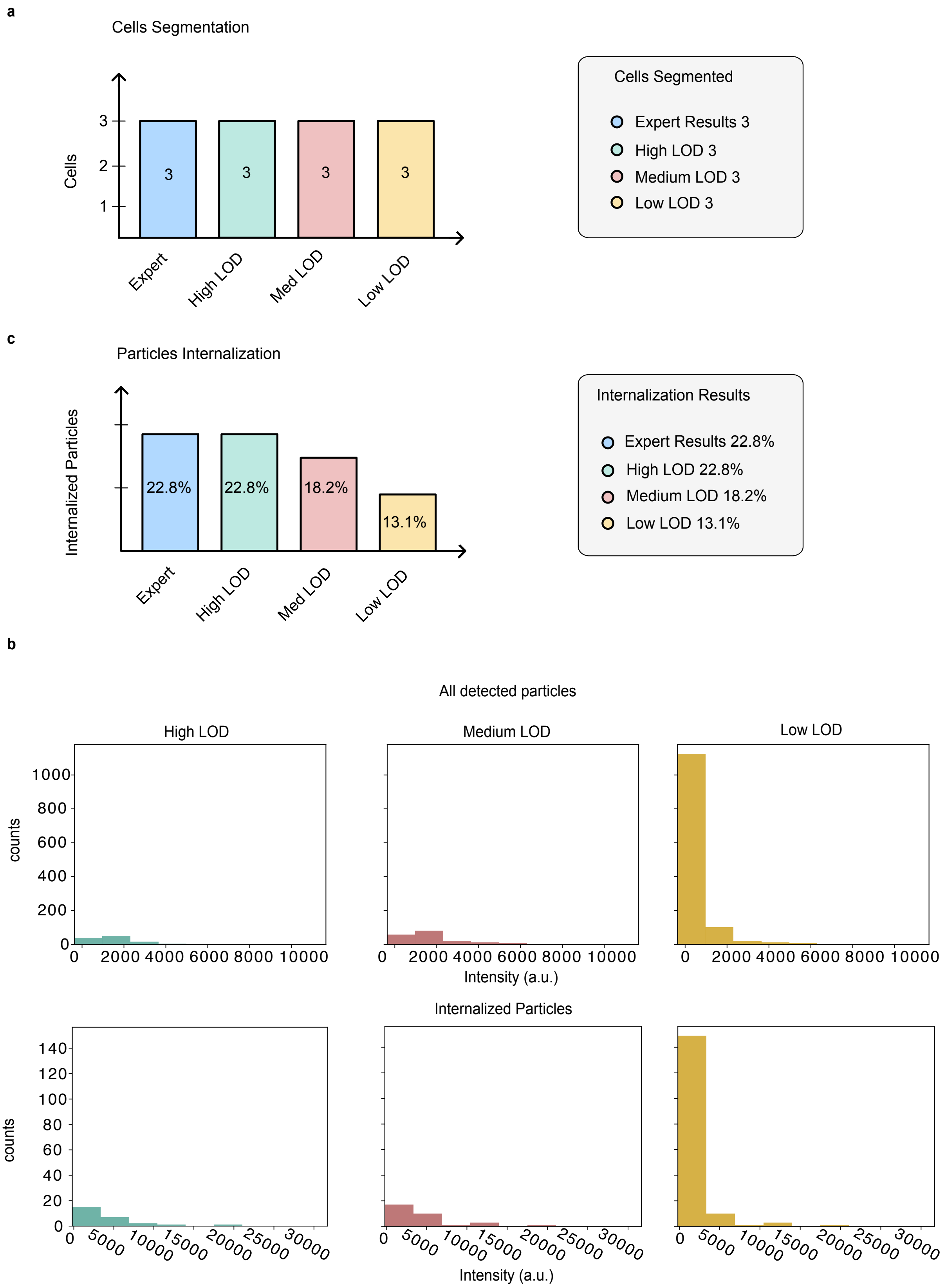


**Supplementary Fig. S6 | LNP internalization outputs across prompt specificity levels.** As this dataset is experimental rather than simulated, the expert-derived analytical reference serves as the benchmark. **(a)** Bar plot of the number of segmented cells for the expert reference and BioBrain outputs obtained under High-, Mid-, and Low-LOD prompts. **(b)** Bar plot of the percentage of internalized particles for the expert reference and BioBrain outputs obtained under High-, Mid-, and Low-LOD prompts. **(c)** Intensity distributions of detected particles (top row) and particles classified as internalized (bottom row) for High-, Mid-, and Low-LOD prompts (columns).

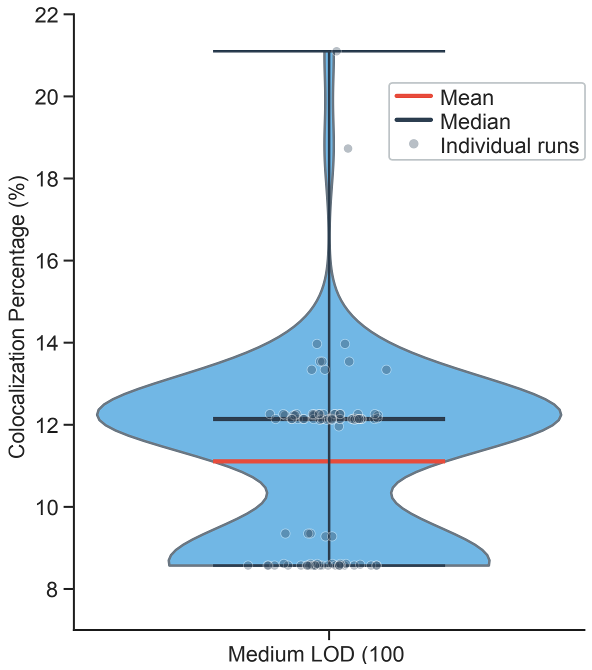


**Supplementary Fig. S7 | Parameter-resolved variability in BioBrain outputs under qualitative prompting.** Violin plot showing the distribution of colocalization percentages at the final endpoint (t120) and inferred detection parameters across 100 independent executions of the Mid-LOD prompt for the cargo–membrane colocalization pipeline. Each run used identical conditions (DeepSeek V3, temperature 0.2, same dataset and file paths). The observed variability reflects a discrete set of parameter combinations selected by the LLM reasoning layer (for example, SNR $\in \{1.5, 2.0\}$ and diameter $\in \{7, 9\}$), each producing a deterministic quantitative output once tools execute, confirming that run-to-run variation is traceable to the inference step rather than to execution instability.

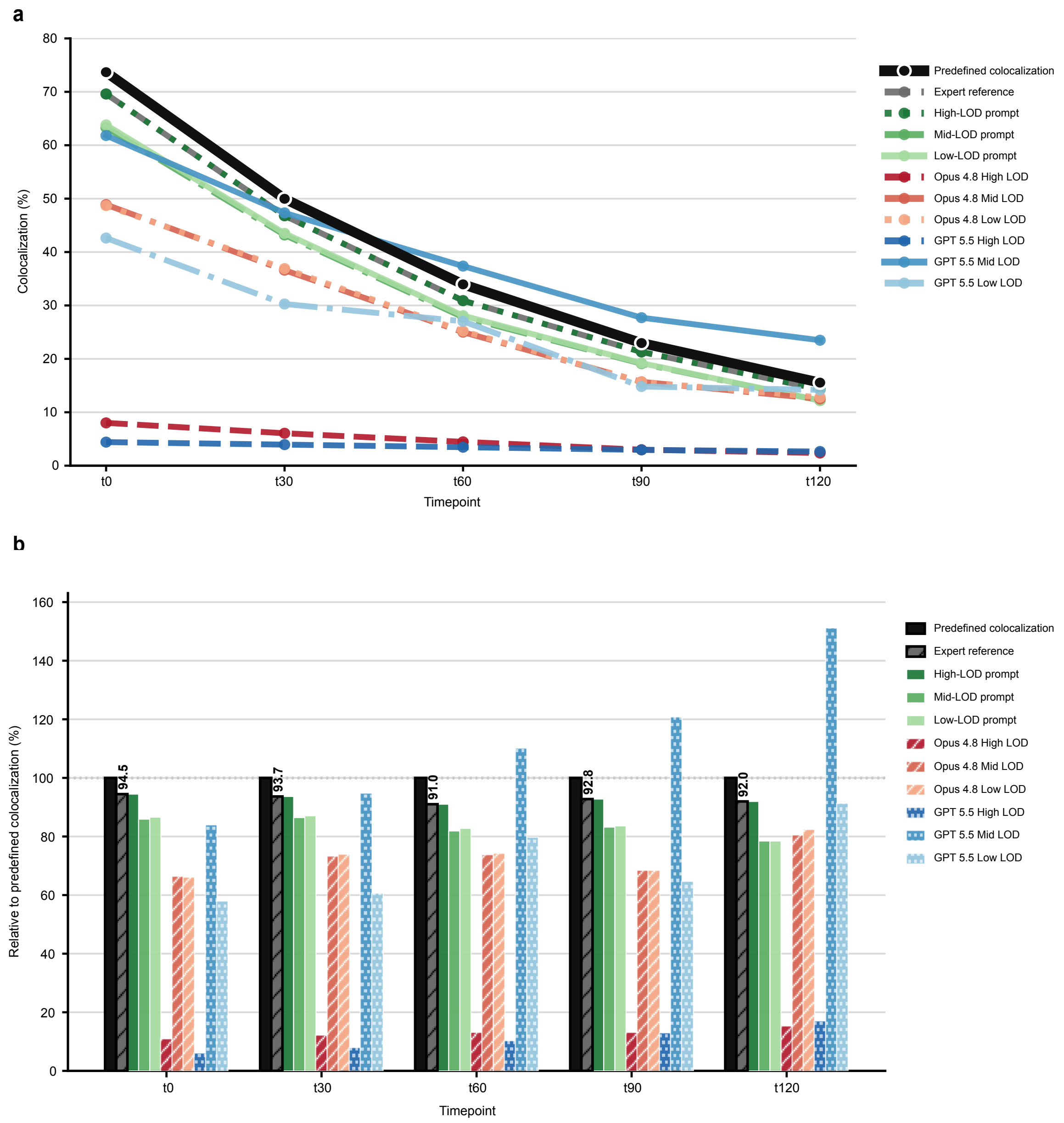


**Supplementary Fig. S8 | Direct, unconstrained LLM analysis of the cargo-membrane colocalization task compared with BioBrain.** The three benchmark prompts (High-, Mid-, and Low-LOD; Supplementary Table S1) were issued directly to two frontier models, Claude Opus 4.8 and GPT-5.5, each given a filesystem path to the same simulated TIRF dataset and allowed to write and execute its own analysis code, without curated tools, schema enforcement, or orchestration. (a) Colocalization fraction at each timepoint (t0–t120) for the simulation ground truth (predefined colocalization), the expert-derived analytical reference, BioBrain outputs under each prompt level, and the directly prompted Opus 4.8 and GPT-5.5 outputs under each prompt level. (b) The same values normalised to the predefined colocalization at each timepoint (100% = predefined ground truth); annotated values above the expert-reference bars give its agreement with the predefined ground truth.

#### Supplementary Tables

**Supplementary Table S1 | Complete prompt texts used for benchmark evaluation.** Exact prompts provided to BioBrain at each specificity level for both the cargo–membrane colocalization and LNP internalization pipelines. High-LOD prompts specify tools and parameters explicitly; Mid-LOD prompts describe intent with qualitative guidance; Low-LOD prompts state only the analytical objective.

##### Cargo–membrane colocalization pipeline

| Prompt level | Prompt text |
|---|---|
| **High-LOD** | <directory_path> contains a time-resolved TIRF microscopy dataset with five discrete timepoints (t0, t30, t60, t90, t120). Each timepoint contains two single-channel TIFF movies: a cargo channel and a membrane channel. Each movie is a 2D time series with dimensions (T, Y, X), consisting of 20 frames of size 512×512 pixels. Each timepoint of the file represents a different subfield of view while the real time dimension can be inferred by the folder name. For each timepoint, convert both cargo and membrane TIFF movies into .npy format. Detect particles independently in the cargo and membrane channels using the Crocker–Grier particle detection method with a particle diameter of 6 pixels and a signal-to-noise ratio of 1.2, treating the data as time-resolved 2D movies. For each timepoint, perform distance-based particle colocalization analysis between cargo and membrane detections using a maximum allowed inter-particle distance of 0.6 pixels. Compute, for each timepoint, the number of and percentage of colocalized particles. Save all intermediate and final outputs—including converted movies, particle detections, colocalization results, and summary statistics—in a structured results directory with distinct filenames. Return the cargo release trend in % per timepoint. |
| **Mid-LOD** | <folder_path> contains a time-resolved microscopy experiment with multiple timepoints. At each timepoint, there are two synchronized movies: one showing intracellular cargo and one showing the cell membrane. For each timepoint, convert the movies into a numerical array format, detect particles in both channels using a particle size appropriate for small vesicular objects and a slightly higher than the default signal-to-noise threshold, and identify cargo particles that are spatially associated with membrane signal within a short interaction distance. Quantify cargo–membrane colocalization at each timepoint and determine cargo release over time. Save all intermediate outputs and summary results in <folder_path> |
| **Low-LOD** | <folder_path> Contains cargo membrane data. What is the percentage of colocalized particles per timepoint? What is the release rate? |

##### LNP internalization pipeline

| Prompt level | Prompt text |
|---|---|
| **High-LOD** | <file_path> is a 3D microscopy volume with axes ordered as (C, Z, Y, X), containing two channels and no time dimension.Channel 0 contains lipid nanoparticle (LNP) signal; channel 1 contains membrane-stained Jurkat cells. Segment cells in 3D using Cellpose (cyto model) on channel 1 with a minimum cell size of 25 voxels and a minimum diameter of 20 voxels. |

| Prompt level | Prompt text |
|---|---|
| | Detect LNP particles in channel 0 using the Crocker–Grier detection method with a signal-to-noise ratio of 1.7 and a particle diameter of 7 voxels. Determine internalization by assigning particles to cells if their centroids lie within cell mask voxels, and compute per-particle internalization status, per-cell counts, and global summary statistics.Save all intermediate and final results in this folder <folder_path>. |
| **Mid-LOD** | <file_path> is a 3D microscopy image with two channels, first one showing LNPs and second one showing cells. The dataset consists of a single 3D volume without a time dimension. I want to quantify lipid nanoparticle internalization into cells.Segment individual cells in 3D using the cell channel with Cellpose and the cyto model. Detect nanoparticle-like objects in the nanoparticle channel, use crocker grier method. The image is a noisy so use a slightly higher SNR than the default. Determine which detected nanoparticles are located inside cells. Assign internalized nanoparticles to individual cells. Compute per-cell internalization statistics and overall summary measurements.Save all intermediate and final results here <folder_path>. |
| **Low-LOD** | <file_path> is a two-channel 3D microscopy image of cells and lipid nanoparticles.I want to quantify how many nanoparticles are internalized into cells. Save all intermediate and final results here <folder_path> |

**Supplementary Table S2 | BioBrain system configuration summary.** Key implementation parameters for all experiments reported in this study.

| **Parameter** | **Value** |
|---|---|
| Implementation language | Python |
| Agent framework | Google Agent Development Kit (ADK) |
| LLM (all agents) | DeepSeek V3 (chat, non-reasoning) |
| Sampling temperature | 0.2 |
| MCP implementation | FastMCP Python package |
| MCP transport protocol | Standard I/O (stdio) |
| Tool schema format | JSON Schema (auto-generated from Python type hints) |
| Agent memory | In-session conversation history (no cross-session persistence) |
| Hardware (TIRF pipeline) | MacBook Pro, Apple M3 Pro |
| Hardware (LNP pipeline) | Workstation with NVIDIA RTX 5000 GPU |
| Execution time (colocalization) | ≈6–7 min |
| Execution time (internalization) | ≈8–9 min (GPU workstation) |
| Source code availability | Not currently public |

**Supplementary Table S3 | BioBrain sub-agents and associated tool categories.** Each sub-agent operates within a defined analytical domain and has access to a curated set of deterministic tools. Tools within each category may include multiple specific implementations. All tools are exposed through the Model Context Protocol with explicit typed input–output schemas.

| **Sub-agent** | **Analytical domain** | **Tool categories** | **Tool origin** |
|---|---|---|---|
| Detection Agent | Particle localisation | Crocker–Grier detection; SNR-based filtering; coordinate extraction | In-house (gpu_tracking) |
| Segmentation Agent | Cell segmentation | 3D instance segmentation; mask generation; minimum size filtering | External (Cellpose 3/4) |
| Colocalization Agent | Spatial association | Distance-based colocalization; per-frame association; temporal aggregation | In-house |
| Internalization Agent | Spatial classification | Centroid-in-mask classification; per-cell counting; summary statistics | In-house |
| File Conversion Agent | Format handling | TIFF–NumPy conversion; OME-TIFF parsing; channel separation | In-house |
| Drift Correction Agent | Registration | Cross-correlation drift estimation; frame-wise coordinate correction | In-house |
| Visualisation Agent | Plotting and rendering | Bar plots; distributions; time-series curves; volume renderings | In-house |
| Tool Integrator Agent | System extensibility | AST analysis; MCP wrapper generation; dependency installation; validation | In-house |

*Note: The sub-agents and tool categories listed above reflect the configuration used for the benchmarks reported in this study. Additional sub-agents and tools can be added to BioBrain at any time through the Tool Integrator Agent or manual registration, and the set of available tools will evolve as the framework is extended to new analytical domains.*
